# A-eye: Automated 3D Segmentation of Healthy Human Eye and Orbit Structures and Axial Length Extraction

**DOI:** 10.1101/2025.05.05.652187

**Authors:** Jaime Barranco, Adrian Luyken, Oliver Stachs, Sönke Langner, Benedetta Franceschiello, Meritxell Bach Cuadra

## Abstract

This study addresses the need for accurate 3D segmentation of the human eye and orbit from MRI to improve ophthalmic diagnostics. Past efforts focused on small sample sizes and varied imaging methods. Here, two techniques (atlas-based registration and supervised deep learning) are tested for automated segmentation on a large T1-weighted MRI dataset. Results show accurate segmentations of the lens, globe, optic nerve, rectus muscles, and fat. Additionally, the study automates the estimation of axial length, a key biomarker.

## 1. Introduction

MRI of the eye (MR-Eye) [1, 2, 3, 4, 5] is raising interest for comprehensive understanding and early disease interception due to its non-invasive and penetration characteristics, able to provide 3D images of the complete eye [6, 7, 8, 9]. MR-Eye can provide key ophthalmic biomarkers: axial length (AL), useful for refractive errors, myopia, hyperopia, glaucoma, and retinal detachment; and volumetry, useful for eye growth abnormalities, glaucoma, macular degeneration, and orbital tumors. It can also advance the development of 3D anatomical models which are helpful for patient-specific treatment planning [9]. To that end, robust and accurate methods for delineation of the eye structures (e.g. lens, vitreous humor, and optic nerve) play a central role.

Pioneer semiautomated methods relied on parametric shape models (spheres and ellipsoids) [10, 11, 12]. However, these deterministic methods based on pre-defined geometry lacked stochastic modelling (e.g. from image intensity, or shape variations) thus capturing anatomical variability. To tackle this, machine learning active shape models (ASM) optimized a robust solution to fit each shape to structures via statistically driven deformations both on healthy eye structures and tumors [13, 14]. Deep learning methods recently appeared in MR-Eye segmentation as well, using a 2D/3D U-Net architectures [15, 16, 17]. Other approaches were combinations of ASM and U-Net [18, 19] and clustering techniques [20]. All these methods are though generally focused on the segmentation of few eye (non-orbit) structures (lens, globe, sclera, cornea, and not so common, optic nerve, only in [14, 15]). Thus, important orbit structures such as rectus muscles or fat, are yet unexplored, jeopardizing the construction of a comprehensive model of the eye and orbit. Moreover, despite methodological efforts (e.g. cross-validation), previous works did not count on a big cohort of manually annotated healthy subjects (sample sizes were limited to 24 to 30 subjects) that would gather properly their anatomical variability for the validation of the results. Finally, most of them (except for [13, 14]) relied on the availability of multi-contrast MRI setting. In this work we present a comprehensive 3D segmentation of the healthy adult human eye and orbit, including lens, globe (vitreous humor), optic nerve, rectus muscles, and fat. We evaluate two different techniques, namely atlas-based segmentation and supervised deep learning (nnU-Net [21]). We quantitatively assess the two methods using a cohort of manually segmented 74 subjects on T1-weighted (T1w). We further explore the application of the two methods on a large-scale cohort of 1157 non-labeled subjects, analyzing eye biometry through automated AL measurement, and provide large-scale automated extraction benchmarks on this ocular MR-Eye biomarker for the first time.

## 2. Materials and Methods

### 2.1. Dataset

Our cohort was originally acquired within the Study of Health in Pomerania (SHIP) [22] and reused in the context of this study. We took 1192/1926 (after quality control) healthy subjects (56±13 y.o., age range 28-89) that underwent whole-body MRI on a 1.5T scanner Magnetom Avanto (Siemens Medical Solutions, Erlangen, Germany) without contrast agent. T1w images of the head were acquired using a 12-channel head coil, voxel dimension 1 mm3, TR=1900 ms, TI=1100 ms, TE=3.37 ms. During the MRI examination, subjects rested their eyes naturally without specific guidelines for viewing or eyelid position. All participants gave informed written consent. The study was approved by the Medical Ethics Committee of the University of Greifswald and followed the Declaration of Helsinki. All data of the study participants were accessed from an anonymized database.

Manual segmentations on a total of 74/1192 subjects were done, using ITK Snap software [23], by two readers independently: one senior (20 years of experience) and one junior (1y). The senior one double checked the annotation by the junior and corrected them if needed. These manual annotations included 9 region-of-interest (ROIs) for the right eye: lens, globe, optic nerve, intraconal and extraconal fats, and the four rectus muscles (lateral, medial, inferior, and superior), see Figure 1B.

**Figure 1:**
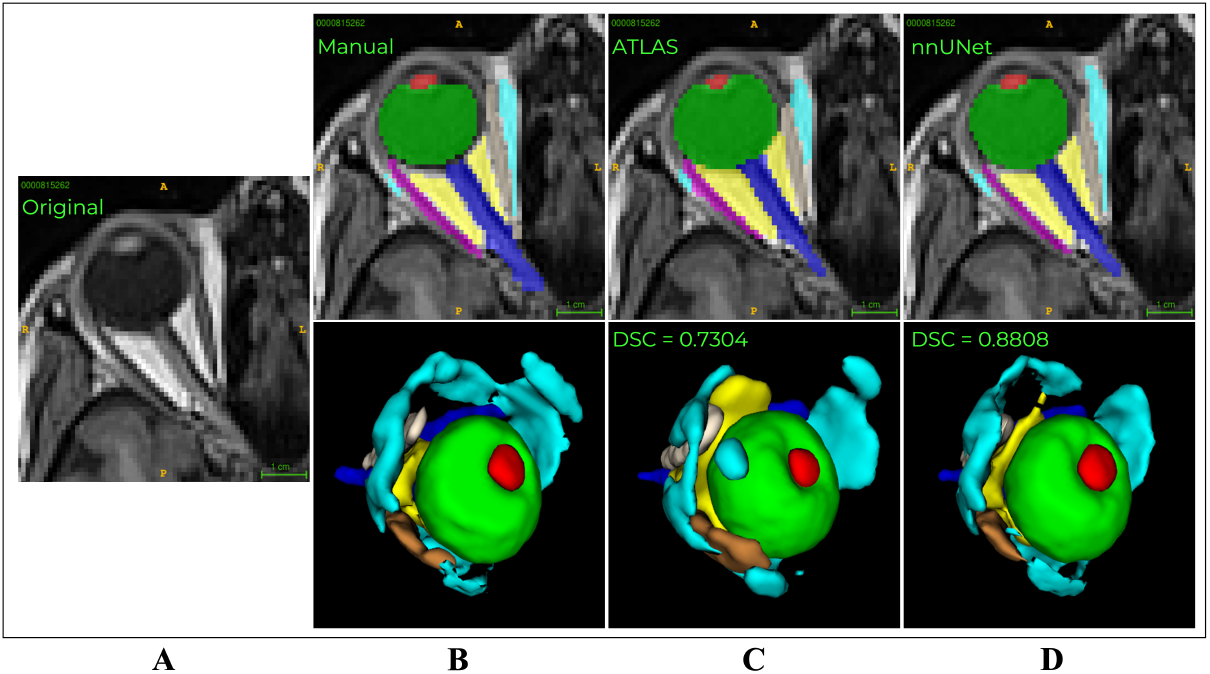
Visual comparison of the different segmentation methods. (A) Original T1w image. (B) Manual segmentation on 9 ROI: lens (red), globe (green), optic nerve (dark blue), intraconal fat (yellow), extraconal fat (cyan), lateral rectus muscle (pink), medial rectus muscle (ivory), inferior rectus muscle (blue), and superior rectus muscle (brown). (C) Atlas-based segmentation. (D) nnU-Net segmentation. We provide preliminary DSC for both automated segmentation methods compared to the manual segmentation (ground truth).

### 2.2. Automated Segmentation Methods

#### Atlas-based segmentation

To the best of our knowledge, atlas-based segmentation was not yet explored in MR-Eye segmentation. However, it is a well established segmentation technique in medical image analysis in general and brain imaging [24, 25]. Atlas-based segmentation relies on voxel-wise image registration of a custom template into the T1w image of the subjects. We used ANTs [26] tool for image registration. Custom template construction: among the manually annotated subjects (N=35/74) (i), the five with highest cross-correlation score after nonlinear registration onto Colin27 [27] were selected (2 females and 3 males); (ii) we then generated a group-wise template T1w map; (iii) we projected label maps of each subject onto the template space using the nonlinear transform in (ii); (iv) we generated the atlas labels by voxel-wise majority voting. Segmentation: (i) first three-stage registration (rigid, affine, and deformable SyN) to determine the bounding box containing the eye (see Figure 1A); (ii) second three-stage registration (rigid, affine, and deformable SyN) within the bounding box; (iii) the atlas’ labels are projected back into the individuals’ spaces. We used a 11th Gen Intel® Core™ i9-11900K × 16 processor with 64GB of RAM. The time spent constructing the template with 5 subjects was about 2.5h, the time of segmentation per subject (considering the whole process) was around 45s, counting a total of around 15h 07m 30s for the “non-trained” cohort of 1157 subjects.

#### Supervised Deep Learning segmentation (nnU-Net)

NnU-Net [21] is the state-of-the-art supervised deep learning-based segmentation approach in which data augmentation is extensively used and the hyperparameters are automatically optimized. It has never been evaluated for MR-Eye, but with Optical Coherence Tomography (OCT) [28]). We split the manual annotated dataset (N=74) into 35 for training (with 4 for evaluation) and 39 for testing (43 in total). Default nnU-Net hyperparameters used: initial learning rate 0.001 with no scheduler; batch size 2; optimizer SGD with momentum 0.99, weight decay 3e-05; dice loss function; data augmentation such as elastic deformation, scaling, rotation; patch size [128, 160, 112]; random parameters initialization; validation strategy cross validation (5 folds); no postprocessing after inference; maximum number of epochs 1000, with an elapsed time of around 140s to 170s per epoch; number of classes 10 (9 ROIs plus background); GPU RTX2080 and RTX3090 (the first available in the cluster), 10 CPUs per task (fold), RAM 64GB, ran in HPC (High Power Computing) SLURM-based cluster, through Docker accessed by Singularity; PyTorch, Python 3.8. The total training time for the five folds was around 208h 20m. The inference process for one image takes about 1 minute and for the whole non-trained dataset (1157 subjects), 66185.53s (18h 23m 05s) with GPU RTX 3060Ti.

### 2.3. Automated Axial Length Extraction from MR-Eye

We developed an algorithm to automatically extract the AL (as defined in [22] and illustrated in Figure 3A). The method inputs both the automated segmented labels and T1w images. First, we computed the centroid of the lens and an orthogonal line to the globe in the 2D axial view, measuring the distance between the extreme intersection points of this line with the lens and the globe. Additionally, we calculated two extra distances: from the globe to the intraconal fat boundary, and from the lens boundary to the cornea. To address the lack of cornea segmentation, we determined the second distance by detecting low-intensity voxels (as the cornea appears black in T1w images) and identifying a significant intensity increase with the help of Sobel filter. This process was applied to multiple slices from the same subject where required segmented structures to compute the measurement were present (lens, globe, optic nerve and intraconal fat). If that was not the case, the AL would be saved as 0 mm. The final AL for a subject was determined by selecting the slice with the highest lens-to-optic nerve ratio (and whose AL was different from 0), indicating the best alignment of these structures to compute the measurement.

### 2.4. Evaluation

#### Segmentation Quality Metrics

To adequately assess the performance of the two segmentation methods, we computed similarity and error metrics between the ground truth (manual segmentations) and the methods’ outputs. Based on [29] we chose Dice Similarity Coefficient (DSC), Hausdorff distance (HD), and volume difference (VD), as they are appropriate metrics to evaluate semantic segmentation of biomedical images.

#### Statistical Analysis

We assessed the significance and effect of differences between methods for each structure and similarity metric. Following the structure in [30], we first tested normality using the Shapiro-Wilk test. Depending on normality, we conducted pair-wise comparisons using either the Wilcoxon test or Student’s t test. A p < 0.05 was considered significant after adjusting for multiple comparisons with the Benjamini-Hochberg’s FDR (q = 0.05). We reported Cohen’s d values to measure effect sizes, with d < 0.5, d > 0.5, and d > 0.8 indicating low, medium, and strong effects, respectively. Mean and confidence intervals (α = 0.95) for Cohen’s d were computed over 5,000 bootstrap samples, selecting 80% of subjects randomly in each iteration.

## 3. Results

Figure 1 presents an example of visual 3D segmented results from the atlas-based and the nnU-Net, as well as the manual reference one.

### Nn-UNet [21] surpassed atlas-based for MR-Eye segmentation tasks

Figure 2 presents boxplots of the results on the 43 pure testing subjects for both methods, and median values with their standard deviations of these results are presented in Table 1. Nn-UNet overall produced better results for the three similarity metrics as compared to the ground truth. Despite the remarkable differences between both methods for DSC and HD, they obtain similar results in terms of VD. For both methods, worse values were encountered in areas more anatomically variable, like the extraconal fat, and smaller structures, i.e. the lens.

**Table 1.** Similarity metrics’ median and standard deviation values for both methods per structure.

| Structure | DSC(nnU-Net, atlas) | HD (nnU-Net, atlas) | VD (nnU-Net, atlas) |
| --- | --- | --- | --- |
| Average | 0.81±0.07, 0.68±0.05 | 0.38±1.3, 0.53±0.26 | 0.15±0.14, 0.01±0.13 |
| Lens | 0.86±0.11, 0.60±0.16 | 0.13±3.56, 0.43±0.87 | 0.16±0.24, -0.59±0.26 |
| Globe | 0.94±0.03, 0.91±0.03 | 0.07±0.05, 0.10±0.09 | 0.06±0.08, 0.06±0.1 |
| Optic nerve | 0.80±0.07, 0.69±0.08 | 0.44±0.26, 0.49±0.22 | -0.02±0.28, 0.13±0.28 |
| Intraconal fat | 0.76±0.11, 0.71±0.06 | 0.40±0.36, 0.43±0.28 | 0.32±0.27, 0.21±0.26 |
| Extraconal fat | 0.70±0.17, 0.58±0.09 | 0.62±2.78, 0.93±0.72 | 0.40±0.32, 0.03±0.33 |
| Lateral RM | 0.82±0.14, 0.70±0.1 | 0.25±0.22, 0.39±0.26 | 0.17±0.25, 0.03±0.27 |
| Medial RM | 0.85±0.09, 0.72±0.07 | 0.19±0.17, 0.40±0.18 | 0.09±0.2, 0.08±0.22 |
| Inferior RM | 0.85±0.08, 0.73±0.06 | 0.21±0.28, 0.38±0.22 | 0.07±0.19, 0.10±0.2 |
| Superior RM | 0.77±0.15, 0.59±0.1 | 0.41±6.97, 0.68±0.64 | 0.18±0.31, 0.21±0.34 |

**Figure 2:**
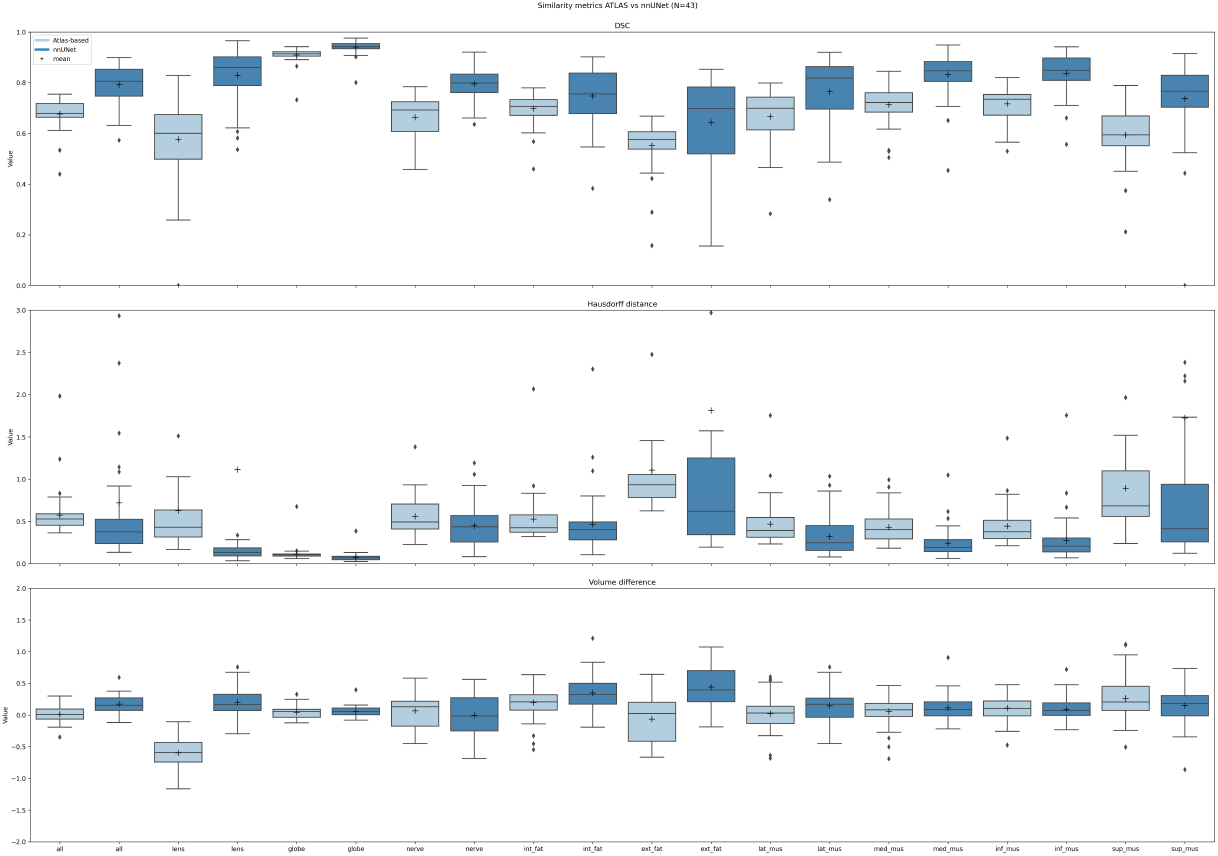
Similarity metrics boxplots computed for both automated segmentation methods on 43 subjects. On the y-axis we have the similarity metric scale (three plots, from top to bottom DSC, HD, VD), and on the x-axis we have the eye-structure (in the following order: all, lens, globe, optic nerve, int. fat, ext. fat, lateral RM, medial RM, inferior RM, and superior RM) per method (light blue for atlas-based, and dark blue for nnU-Net).

**Figure 3:**
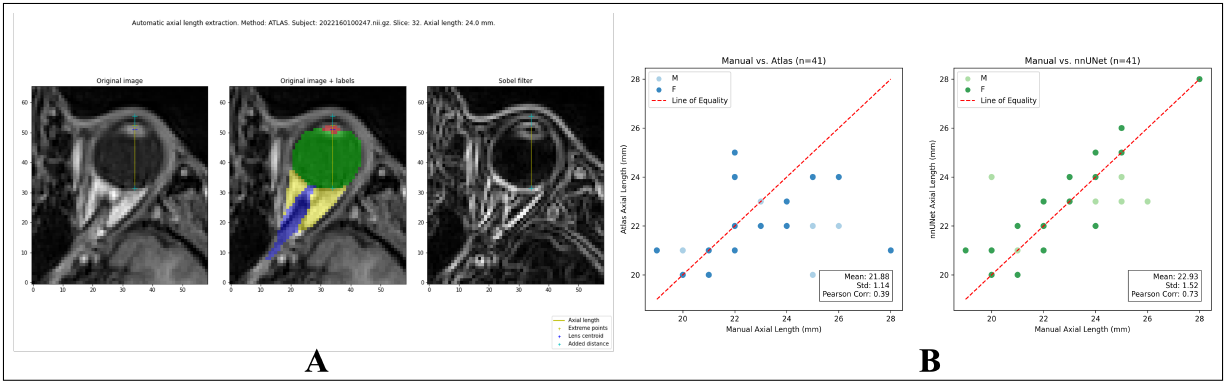
Axial length (posterior surface of the cornea to the posterior pole of the ocular bulb, at the boundary with orbital fat) per method grouped by sex. (A) Example of AL extraction in MRI: on the left, manual extraction from [22]; in the middle, automatic extraction using the segmented structures and the T1w image; on the right, Sobel-filtered image visual aid. (B) Correlation curves and coefficient on N=41 with respect to the subject-wise manual annotations, which serve as ground truth.

### Overlap and surface distance metrics from both methods are statistically significantly different

In Table 2, we present the significance and effect (magnitude) of the differences between both methods. They are statistically different specially in DSC, having strong and medium effect across structures. In HD, while having low and medium effects, the p values showed a significant difference (p<0.05), whereas in VD, p values showed no significant difference except for lens and fat, with strong effects.

**Table 2.**
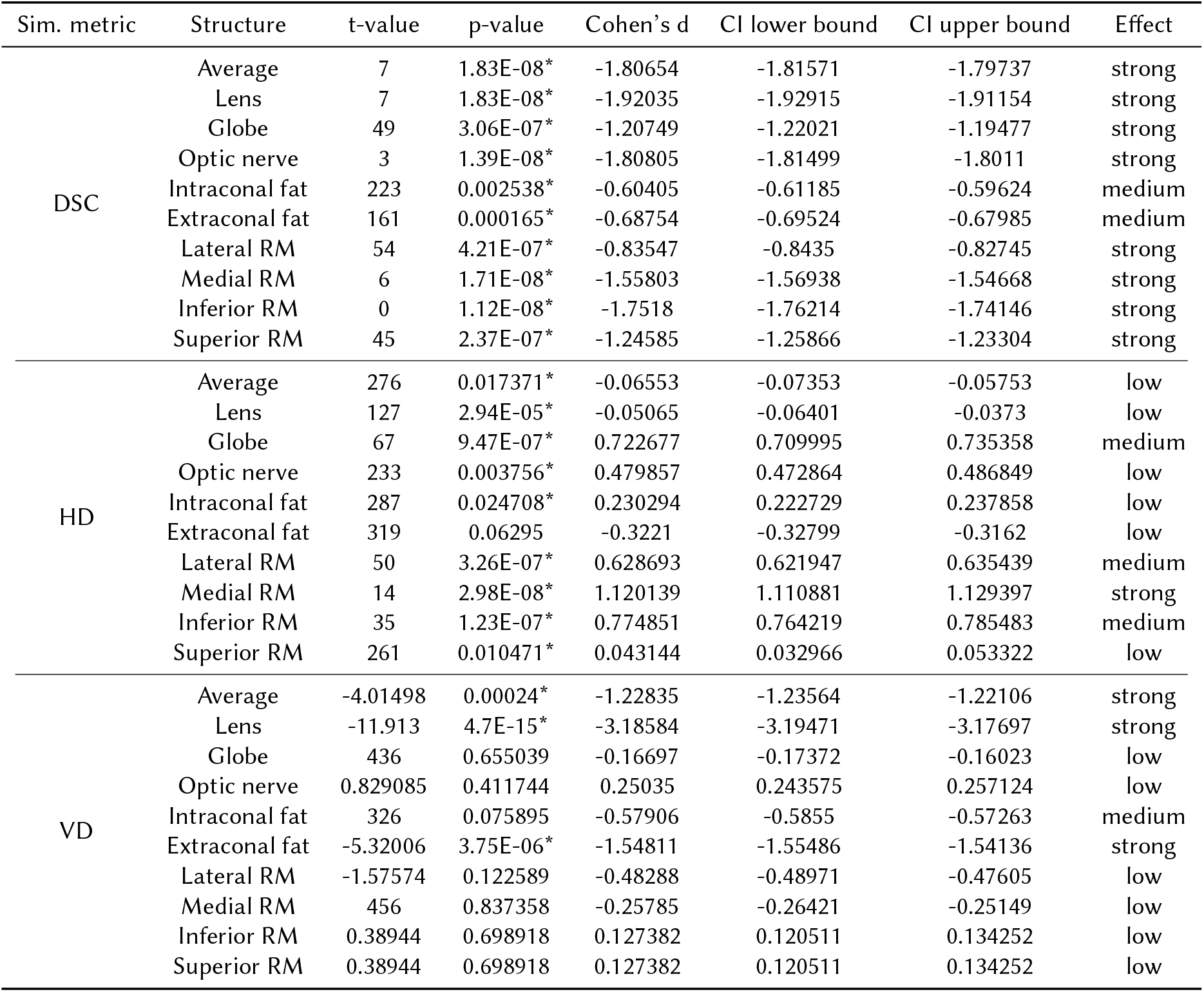
Significance and effect of the differences in atlas-based and nnU-Net methods regarding similarity metrics per eye structure. The most impactful difference is in DSC, having strong and medium effect across structures. In HD, while having low and medium effects, the p values showed a significant difference (p<0.05, marked with *), whereas in VD, p values showed no significant difference except for lens and fat, with strong effects.

### Automated measurements of axial length from MRI are in line with the reference manual measures for both methods, but closer to nnU-Net

Figure 3B shows the correlation plots of the extracted values per method, grouped by sex, as in [22], on the manually annotated cohort of 43 testing subjects, resulting in 41 after removing zero values. The mean AL from nnU-Net segmentation was closer to the ground truth values which were extracted using the same extraction method on the manual segmentations. For the large-scale cohort of 1157 subjects, the mean values and standard deviations, after removing zero values for both methods, show closer values to the manual reference for the nnU-Net case as well:

- nnU-Net: 23.8±1.7 mm (519 M) and 22.9±1.6 mm (522 F)
- Atlas-based: 22.3±1.2 mm (519 M) and 21.7±1.3 mm (522 F)
- Previous studies [22]: 23.4±0.8 mm (1059 M) and 22.8±0.9 mm (867 F)

## 4. Discussion

In this work, we have introduced an evaluation of two known automated 3D segmentation methods, atlas-based and nnU-Net, in the new application field of eye and orbit structures in MRI. These techniques enabled us to analyze eye anatomy by extracting axial length measurements from a large-scale cohort, thereby establishing benchmarks for automated AL measurements.

Supervised deep learning segmentation (nnU-Net) of eye structures surpassed atlas-based method in 43 pure testing subjects, based on three complementary evaluation metrics (DSC, HD, VD) and axial length extractions. Based on these quantitative results, nnU-Net should be preferred for MR-Eye segmentation tasks. However, our work is still preliminary, evaluated only in-domain and further analysis in out-of-domain testing would be required.

Further developments will include the automated extraction of other important ophthalmic biomarkers and explore their possible correlation with other biomarkers like body mass index [22]. In the context of large-scale segmentation, it would also be important to include uncertainty quantification of automated predictions [31]. Also, it would be very relevant to have a comprehensive qualitative evaluation of the resulting segmentations and biomarkers by clinicians. Finally, our large-scale study builds towards a ready-to-use solution to promote the adoption of accurate MR-Eye segmentation and its further applications by clinicians and researchers, boosting clinical dissemination and translation of MR-Eye.

## Acknowledgments

We would like to thank the following contributors: Hamza Kebiri^1^, Philipp Stachs^7^, Pedro M. Gordaliza^1^, Oscar Esteban^2^, Yasser Aleman^2^, Raphael Sznitman^8*7*^*KIT*, ^*8*^*ARTORG*

**Conceptualization:** MBC, BF, SL, OS **Dataset:** SL, OS, PS, AL **Methodology—Atlas registration:** JB, OE, YA, MBC **Methodology—Deep Learning, nnU-Net:** JB, HK, PMG, MBC **Methodology—QA/QC:** JB, OE, BF, MBC **Methodology—Biomarkers extraction:** JB, BF, MBC **Methodology—Clinical relevance:** JB, SL, OS, AL, BF, MBC **Methodology—Atlas of the eye and maps of the labels:** JB, YA, BF, MBC **Methodology—Statistics:** JB, YA, BF, MBC **Supervision:** MBC, BF, SL, OS **Writing—review & editing:** JB, MBC, BF, SL, OS, AL, OE, HK, PMG

## A. Appendix: tables

## References

[1] T. Georgouli, T. James, S. Tanner, D. Shelley, M. Nelson, B. Chang, O. Backhouse, D. McGonagle, High-Resolution Microscopy Coil MR-Eye, Eye 22 (2008) 994–996. URL: https://www.nature.com/articles/6702755. doi:10.1038/sj.eye.6702755.

[2] I. Tsiapa, M. K. Tsilimbaris, E. Papadaki, P. Bouziotis, I. G. Pallikaris, A. H. Karantanas, T. G. Maris, High resolution MR eye protocol optimization: Comparison between 3D-CISS, 3D-PSIF and 3D-VIBE sequences, Physica Medica 31 (2015) 774–780. URL: https://linkinghub.elsevier.com/retrieve/pii/S112017971500071X. doi:10.1016/j.ejmp.2015.03.009.

[3] N. Dobbs, M. Budak, R. White, I. Zealley, MR-Eye: High-Resolution Microscopy Coil MRI for the Assessment of the Orbit and Periorbital Structures, Part 1: Technique and Anatomy, American Journal of Neuroradiology 41 (2020) 947–950. URL: http://www.ajnr.org/lookup/doi/10.3174/ajnr.A6495. doi:10.3174/ajnr.A6495.

[4] E. Fleury, P. Trnková, E. Erdal, M. Hassan, B. Stoel, M. Jaarma-Coes, G. Luyten, J. Herault, A. Webb, J. Beenakker, J. Pignol, M. Hoogeman, Three-dimensional MRI-based treatment planning approach for non-invasive ocular proton therapy, Medical Physics 48 (2021) 1315–1326. URL: https://aapm.onlinelibrary.wiley.com/doi/10.1002/mp.14665. doi:10.1002/mp.14665.

[5] R. K. Glarin, B. N. Nguyen, J. O. Cleary, S. C. Kolbe, R. J. Ordidge, B. V. Bui, A. M. McKendrick, B. A. Moffat, MR-EYE: High-Resolution MRI of the Human Eye and Orbit at Ultrahigh Field (7T), Magnetic Resonance Imaging Clinics of North America 29 (2021) 103–116. URL: https://linkinghub.elsevier.com/retrieve/pii/S1064968920300696. doi:10.1016/j.mric.2020.09.004.

[6] K. A. Townsend, G. Wollstein, J. S. Schuman, Clinical application of MRI in ophthalmology, NMR in Biomedicine 21 (2008) 997–1002. URL: https://analyticalsciencejournals.onlinelibrary.wiley.com/doi/10.1002/nbm.1247. doi:10.1002/nbm.1247.

[7] L. Fanea, A. J. Fagan, Review: Magnetic resonance imaging techniques in ophthalmology, Molecular Vision 18 (2012) 2538–2560. URL: https://www.ncbi.nlm.nih.gov/pmc/articles/PMC3482169/.

[8] T. Q. Duong, Magnetic resonance imaging of the retina: From mice to men, Magnetic Resonance in Medicine 71 (2014) 1526–1530. URL: https://onlinelibrary.wiley.com/doi/10.1002/mrm.24797. doi:10.1002/mrm.24797.

[9] T. Niendorf, J.-W. M. Beenakker, S. Langner, K. Erb-Eigner, M. Bach Cuadra, E. Beller, J. M. Millward, T. M. Niendorf, O. Stachs, Ophthalmic Magnetic Resonance Imaging: Where Are We (Heading To)?, Current Eye Research 46 (2021) 1251–1270. URL: https://www.tandfonline.com/doi/full/10.1080/02713683.2021.1874021. doi:10.1080/02713683.2021.1874021.

[10] M. Goitein, T. Miller, Planning proton therapy of the eye, Medical Physics 10 (1983) 275–283. doi:10.1118/1.595258.

[11] B. Dobler, R. Bendl, Precise modelling of the eye for proton therapy of intra-ocular tumours, Physics in Medicine and Biology 47 (2002) 593–613. doi:10.1088/0031-9155/47/4/304.

[12] K. D. Singh, N. S. Logan, B. Gilmartin, Three-dimensional modeling of the human eye based on magnetic resonance imaging, Investigative Ophthalmology & Visual Science 47 (2006) 2272–2279. doi:10.1167/iovs.05-0856.

[13] C. Ciller, S. I. De Zanet, M. B. Rüegsegger, A. Pica, R. Sznitman, J.-P. Thiran, P. Maeder, F. L. Munier, J. H. Kowal, M. B. Cuadra, Automatic Segmentation of the Eye in 3D Magnetic Resonance Imaging: A Novel Statistical Shape Model for Treatment Planning of Retinoblastoma, International Journal of Radiation Oncology*Biology*Physics 92 (2015) 794–802. URL: https://linkinghub.elsevier.com/retrieve/pii/S0360301615002990. doi:10.1016/j.ijrobp.2015.02.056.

[14] H.-G. Nguyen, R. Sznitman, P. Maeder, A. Schalenbourg, M. Peroni, J. Hrbacek, D. C. Weber, A. Pica, M. Bach Cuadra, Personalized Anatomic Eye Model From T1-Weighted Volume Interpolated Gradient Echo Magnetic Resonance Imaging of Patients With Uveal Melanoma, International Journal of Radiation Oncology*Biology*Physics 102 (2018) 813–820. URL: https://linkinghub.elsevier.com/retrieve/pii/S0360301618307971. doi:10.1016/j.ijrobp.2018.05.004.

[15] O. Ronneberger, P. Fischer, T. Brox, U-Net: Convolutional Networks for Biomedical Image Segmentation, 2015. URL: http://arxiv.org/abs/1505.04597. doi:10.48550/arXiv.1505.04597, 1505.04597 [cs].

[16] H.-G. Nguyen, A. Pica, P. Maeder, A. Schalenbourg, M. Peroni, J. Hrbacek, D. C. Weber, M. B. Cuadra, R. Sznitman, Ocular Structures Segmentation from Multi-sequences MRI Using 3D Unet with Fully Connected CRFs, in: D. Stoyanov, Z. Taylor, F. Ciompi, Y. Xu, A. Martel, L. Maier-Hein, N. Rajpoot, J. van der Laak, M. Veta, S. McKenna, D. Snead, E. Trucco, M. K. Garvin, X. J. Chen, H. Bogunovic (Eds.), Computational Pathology and Ophthalmic Medical Image Analysis, Lecture Notes in Computer Science, Springer International Publishing, Cham, 2018, pp. 167–175. doi:10.1007/978-3-030-00949-6_20.

[17] V. I. J. Strijbis, C. M. De Bloeme, R. W. Jansen, H. Kebiri, H.-G. Nguyen, M. C. De Jong, A. C. Moll, M. Bach-Cuadra, P. De Graaf, M. D. Steenwijk, Multi-view convolutional neural networks for automated ocular structure and tumor segmentation in retinoblastoma, Scientific Reports 11 (2021) 14590. URL: https://www.nature.com/articles/s41598-021-93905-2. doi:10.1038/s41598-021-93905-2.

[18] C. Ciller, S. De Zanet, K. Kamnitsas, P. Maeder, B. Glocker, F. L. Munier, D. Rueckert, J.-P. Thiran, M. Bach Cuadra, R. Sznitman, Multi-channel MRI segmentation of eye structures and tumors using patient-specific features, PLOS ONE 12 (2017) e0173900. URL: https://dx.plos.org/10.1371/journal.pone.0173900. doi:10.1371/journal.pone.0173900.

[19] H.-G. Nguyen, A. Pica, F. L. Rosa, J. Hrbacek, D. C. Weber, A. Schalenbourg, R. Sznitman, M. B. Cuadra, A novel segmentation framework for uveal melanoma based on magnetic resonance imaging and class activation maps (2019). URL: https://boris.unibe.ch/135253/. doi:10.7892/BORIS.135253.

[20] M. K. Hassan, E. Fleury, D. Shamonin, L. G. Fonk, M. Marinkovic, M. G. Jaarsma-Coes, G. P. Luyten, A. Webb, J.-W. Beenakker, B. Stoel, An Automatic Framework to Create Patient-specific Eye Models From 3D Magnetic Resonance Images for Treatment Selection in Patients With Uveal Melanoma, Advances in Radiation Oncology 6 (2021) 100697. URL: https://linkinghub.elsevier.com/retrieve/pii/S2452109421000555. doi:10.1016/j.adro.2021.100697.

[21] F. Isensee, P. F. Jaeger, S. A. A. Kohl, J. Petersen, K. H. Maier-Hein, nnU-Net: a self-configuring method for deep learning-based biomedical image segmentation, Nature Methods 18 (2021) 203–211. URL: https://www.nature.com/articles/s41592-020-01008-z. doi:10.1038/s41592-020-01008-z.

[22] P. Schmidt, R. Kempin, S. Langner, A. Beule, S. Kindler, T. Koppe, H. Völzke, T. Ittermann, C. Jürgens, F. Tost, Association of anthropometric markers with globe position: A population-based MRI study, PLoS ONE 14 (2019) e0211817. URL: https://www.ncbi.nlm.nih.gov/pmc/articles/PMC6366780/. doi:10.1371/journal.pone.0211817.

[23] P. A. Yushkevich, J. Piven, H. C. Hazlett, R. G. Smith, S. Ho, J. C. Gee, G. Gerig, User-guided 3D active contour segmentation of anatomical structures: Significantly improved efficiency and reliability, NeuroImage 31 (2006) 1116–1128. URL: https://www.sciencedirect.com/science/article/pii/S1053811906000632. doi:10.1016/j.neuroimage.2006.01.015.

[24] M. Bach Cuadra, V. Duay, J.-P. Thiran, Atlas-based Segmentation, in: N. Paragios, J. Duncan, N. Ayache (Eds.), Handbook of Biomedical Imaging: Methodologies and Clinical Research, Springer US, Boston, MA, 2015, pp. 221–244. URL: https://doi.org/10.1007/978-0-387-09749-7_12. doi:10.1007/978-0-387-09749-7_12.

[25] M. Cabezas, A. Oliver, X. Lladó, J. Freixenet, M. Bach Cuadra, A review of atlas-based segmentation for magnetic resonance brain images, Computer Methods and Programs in Biomedicine 104 (2011) e158–e177. URL: https://www.sciencedirect.com/science/article/pii/S0169260711002033. doi:10.1016/j.cmpb.2011.07.015.

[26] B. Avants, N. J. Tustison, G. Song, Advanced Normalization Tools: V1.0, The Insight Journal (2009). URL: https://www.insight-journal.org/browse/publication/681. doi:10.54294/uvnhin.

[27] C. J. Holmes, R. Hoge, L. Collins, R. Woods, A. W. Toga, A. C. Evans, Enhancement of MR Images Using Registration for Signal Averaging, Journal of Computer Assisted Tomography 22 (1998) 324. URL: https://journals.lww.com/jcat/abstract/1998/03000/enhancement_of_mr_images_using_registration_for.32.aspx.

[28] P. Valmaggia, P. Friedli, B. Hörmann, P. Kaiser, H. P. N. Scholl, P. C. Cattin, R. Sandkühler, P. M. Maloca, Feasibility of Automated Segmentation of Pigmented Choroidal Lesions in OCT Data With Deep Learning, Translational Vision Science & Technology 11 (2022) 25. URL: https://tvst.arvojournals.org/article.aspx?articleid=2783676. doi:10.1167/tvst.11.9.25.

[29] L. Maier-Hein, A. Reinke, P. Godau, M. D. Tizabi, F. Buettner, E. Christodoulou, B. Glocker, F. Isensee, J. Kleesiek, M. Kozubek, M. Reyes, M. A. Riegler, M. Wiesenfarth, A. E. Kavur, C. H. Sudre, M. Baumgartner, M. Eisenmann, D. Heckmann-Nötzel, A. T. Rädsch, L. Acion, M. Antonelli, T. Arbel, S. Bakas, A. Benis, M. Blaschko, M. J. Cardoso, V. Cheplygina, B. A. Cimini, G. S. Collins, K. Farahani, L. Ferrer, A. Galdran, B. van Ginneken, R. Haase, D. A. Hashimoto, M. M. Hoffman, M. Huisman, P. Jannin, C. E. Kahn, D. Kainmueller, B. Kainz, A. Karargyris, A. Karthikesalingam, H. Kenngott, F. Kofler, A. Kopp-Schneider, A. Kreshuk, T. Kurc, B. A. Landman, G. Litjens, A. Madani, K. Maier-Hein, A. L. Martel, P. Mattson, E. Meijering, B. Menze, K. G. M. Moons, H. Müller, B. Nichyporuk, F. Nickel, J. Petersen, N. Rajpoot, N. Rieke, J. Saez-Rodriguez, C. I. Sánchez, S. Shetty, M. van Smeden, R. M. Summers, A. A. Taha, A. Tiulpin, S. A. Tsaftaris, B. Van Calster, G. Varoquaux, P. F. Jäger, Metrics reloaded: Recommendations for image analysis validation, 2023. URL: http://arxiv.org/abs/2206.01653. doi:10.48550/arXiv.2206.01653, 2206.01653 [cs].

[30] Y. Alemán-Gómez, E. Najdenovska, T. Roine, M. J. Fartaria, E.J. Canales-Rodríguez, Z. Rovó, P. Hagmann, P. Conus, K. Q. Do, P. Klauser, P. Steullet, P. S. Baumann, M. Bach Cuadra, Partial-volume modeling reveals reduced gray matter in specific thalamic nuclei early in the time course of psychosis and chronic schizophrenia, Human Brain Mapping 41 (2020) 4041–4061. URL: https://www.ncbi.nlm.nih.gov/pmc/articles/PMC7469814/. doi:10.1002/hbm.25108.

[31] B. Lambert, F. Forbes, S. Doyle, H. Dehaene, M. Dojat, Trustworthy clinical AI solutions: A unified review of uncertainty quantification in Deep Learning models for medical image analysis, Artificial Intelligence in Medicine 150 (2024) 102830. URL: https://www.sciencedirect.com/science/article/pii/S0933365724000721. doi:10.1016/j.artmed.2024.102830.

